# Evolutionarily developed connections compromised in schizophrenia

**DOI:** 10.1101/387506

**Authors:** Martijn P. van den Heuvel, Lianne H. Scholtens, Siemon C. de Lange, Rory Pijnenburg, Wiepke Cahn, Neeltje E.M. van Haren, Iris Sommer, Longchuan Li, Todd Preuss, James K. Rilling

## Abstract

The genetic basis and uniquely human character of schizophrenia has led to the notion of human brain evolution to have resulted in vulnerability to the disorder. We examined schizophrenia-related changes in brain connectivity in the context of evolutionary changes in human brain wiring by comparing *in-vivo* neuroimaging data from humans, chimpanzees and macaque monkeys. We find that evolutionary changes in human connectome organization overlap with the pattern of schizophrenia-related changes in brain connectivity, with connections evolutionary enhanced in the human brain showing significantly more involvement in schizophrenia pathology than connections shared between humans and non-human primates (effects shown in three independent patient-control datasets). Our findings suggest that the evolution of brain wiring in support of complex brain function in humans may have come at the cost of an increased vulnerability to brain dysfunction in disease.

## Introduction

Schizophrenia is a severe psychiatric disorder that is characterized by hallucinations, delusions, loss of initiative and general cognitive dysfunction. Its high heritability and cross-cultural ubiquity led to the early hypothesis of a genetic origin for the disorder, an idea that has gained strong support from modern-day large-scale genetic studies (1). Its genetic basis coupled with the notion of the disorder to be present across societies that all originated from the same modern human predecessor but otherwise lived mostly isolated until the last few centuries, under large variation in climate, level of urbanisation, culture, industrialisation etcetera, has led to the suggestion that human brain evolution may have played an important role in vulnerability to the disorder (2).

Crow’s original theory of an evolutionary origin of psychosis (2) is motivated on one hand by the observation of enhanced cerebral asymmetry in humans compared to other primates and on the other hand by empirical observations of reduced corpus callosum connectivity and asymmetrical changes of ventricular size in patients with schizophrenia (e.g. (3) (4)). In addition, Burns further argued that schizophrenia may be the result of the evolution of complex social cognition (5) and accompanying development of interregional connectivity in hominins, rendering the brain vulnerable to alterations with this complex brain wiring.

Schizophrenia has been suggested to be a costly by-product of increasing brain size and accompanied complexity of brain connectivity, with an increase in connectivity leading to heightened vulnerability for the development of brain disorders (6). Indeed, emerging evidence from modern-day studies of the network structure of the human brain –the human connectome– have suggested that schizophrenia may involve a disruption of the brain’s large-scale connectivity architecture (see (7) for review). This collectively leads to the question whether unique evolutionary changes in the human connectome played a role in the emergence of cerebral disconnectivity associated with schizophrenia.

One approach to investigating the evolution of human brain connectivity is to compare human brain organization with that of other living primate species, especially with our closest living primate relative, the chimpanzee. Our two species share a last common African ancestor from about 5-10 million years ago, making chimpanzees an ideal group to study human evolutionary brain development. Comparative primate research suggests that larger brain size is associated with reduced inter-hemispheric and increased intra-hemispheric connectivity (8), and this relative increase in white matter volume compared to total brain size marks an increasing investment of neural resources that enable efficient communication between brain regions. As such, encephalization may involve an increasing dependence of brain function on the effectiveness of evolutionary new connectivity (9).

In line with the aforementioned evolutionary hypothesis of schizophrenia, a selection pressure for effective information transfer between remote brain regions in service of high-level brain function may be paralleled by an increasing risk of disrupted brain function, and with that arguably a higher risk of emergence of human-unique brain disorders. Here, we adopted the framework of *comparative connectomics* (10) –the field that studies commonalities and differences in the topological organization of brain circuits across species– to evaluate the proposed role of brain evolution, and in particular those of hominids, in the emergence of schizophrenia. We compared human brain network organization with that of chimpanzees and show evidence of overlap in the pattern of schizophrenia brain disconnectivity and human evolutionary changes to brain connectivity.

## Results

### Chimpanzee - human network comparison

Chimpanzee (n=22) and human (n=58) connectome maps were reconstructed on the basis of acquired high-resolution T1 and diffusion-weighted MRI data, consisting of 114 cortical regions (DK-114) and all reconstructed cortico-cortical connections between these regions (see methods). Other parcellation schemes (DK-68 and DK-219) and the use of a species homologue atlas (EBB-38, see below) resulted in similar results, see Supporting Materials and robustness analysis below). The chimpanzee and human brain network showed large overlap in their overall connectome layout, with a binary overlap of 94% (p<0.001, Mantel test) and a strong overall correlation in connectivity strength (FA-weighted, r=0.93, p<0.001, Figure 1).

**Figure 1.**
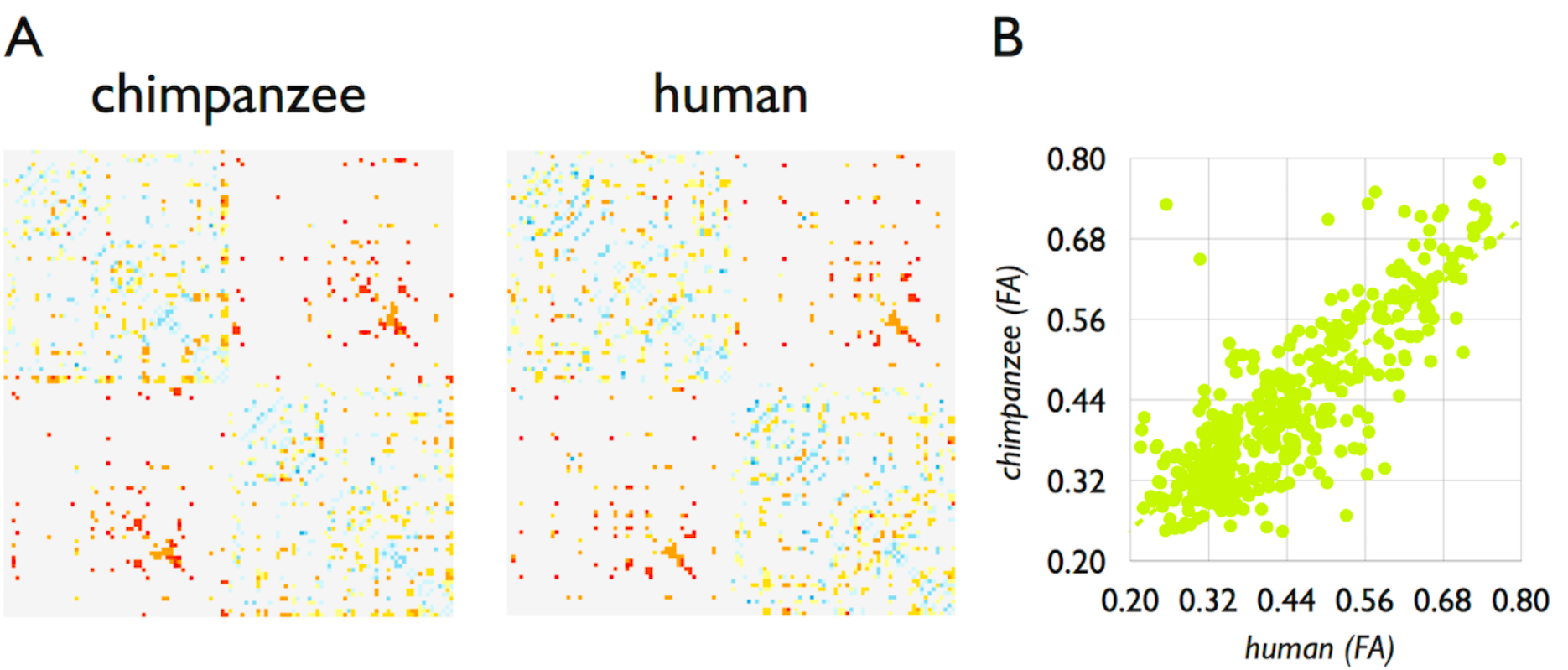
Chimpanzee and Human connectome. **Panel A** shows the group-averaged chimpanzee (n=22, 60% group threshold) and human connectome (n=58, 60% threshold group connectome). Colored values show (normalized) FA values of the chimpanzee and human connectome from low (blue) to high (red). Non-colored pixels represent no connection present. **Panel B** shows the correlation between FA values of all shared connections of the two species (r=0.93, p<0.001).

We next selected *human-specific connections* as the set of cortico-cortical pathways observed in at least 60% of the group of humans and not in any of the chimpanzees. A strict prevalence group threshold of 60% (human) and an exclusion threshold of 0% (chimpanzee) were used to reduce the inclusion of false-positive reconstructions at the group level (11) and to include a strict definition of human-specific tracts (other group thresholds revealed similar findings, see Supplemental Information (SI)). 27 human-specific connections of the reconstructed group-average human connectome were observed (3.5% of total connections). These human-specific connections included cortico-cortical connections connecting frontal, temporal and parietal brain regions, including connections of the posterior bank of the left superior temporal sulcus (bank), left inferior and middle temporal gyri, left posterior inferior parietal cortex, left caudal anterior cingulate cortex and lateral orbitofrontal cortex, ventral part of the right and left precentral gyrus, left pars opercularis left paracentral cortex, left precuneus, right cuneus, right insula, right superior frontal cortex, right medial orbitofrontal and pars orbitalis and right inferior and superior parietal cortex (listed in Supporting Table 1 and shown in Figure 3).

For contrast, we also computed chimpanzee-specific connections (i.e. reconstructed pathways observed in at least 60% of chimpanzees, but not in humans), which yielded only 14 connections (2.3% of the chimpanzee connectome).

Comparison also revealed *human-chimpanzee shared connections*, taken as the set of connections that were observed in at least 60% of both the groups of humans and chimpanzees, and included a total set of 428 connections (56% of the human connectome, 72% of the chimpanzee connectome). To reduce the potential inclusion of false positive tracts (11) and to thus use a strict definition of human-specific and human-chimpanzee shared connections (12) other connections (i.e. tracts observed in less than 60% of the groups and of which one can not be sure whether they represent true positives) were not considered for further analysis.

### Schizophrenia brain disconnectivity

We continued by examining the pattern of brain disconnectivity related to schizophrenia, assessed by edge-wise statistical comparison of connectivity strength (FA-weighted, see methods) across the principal set of n=48 schizophrenia patients and n=43 matched healthy controls (Supporting Table 2). Strongest patient-control differences were observed around connections of the ventral part of the posterior cingulate cortex (isthmus), inferior and superior parietal and inferior and superior temporal cortex, frontal pole, insular cortex, rostral middle frontal cortex, precuneus, supramarginal gyrus and post and precentral gyrus (Network Based Statistics p=0.024, Figure 2); findings consistent with previous schizophrenia connectome studies (12-14).

**Figure 2.**
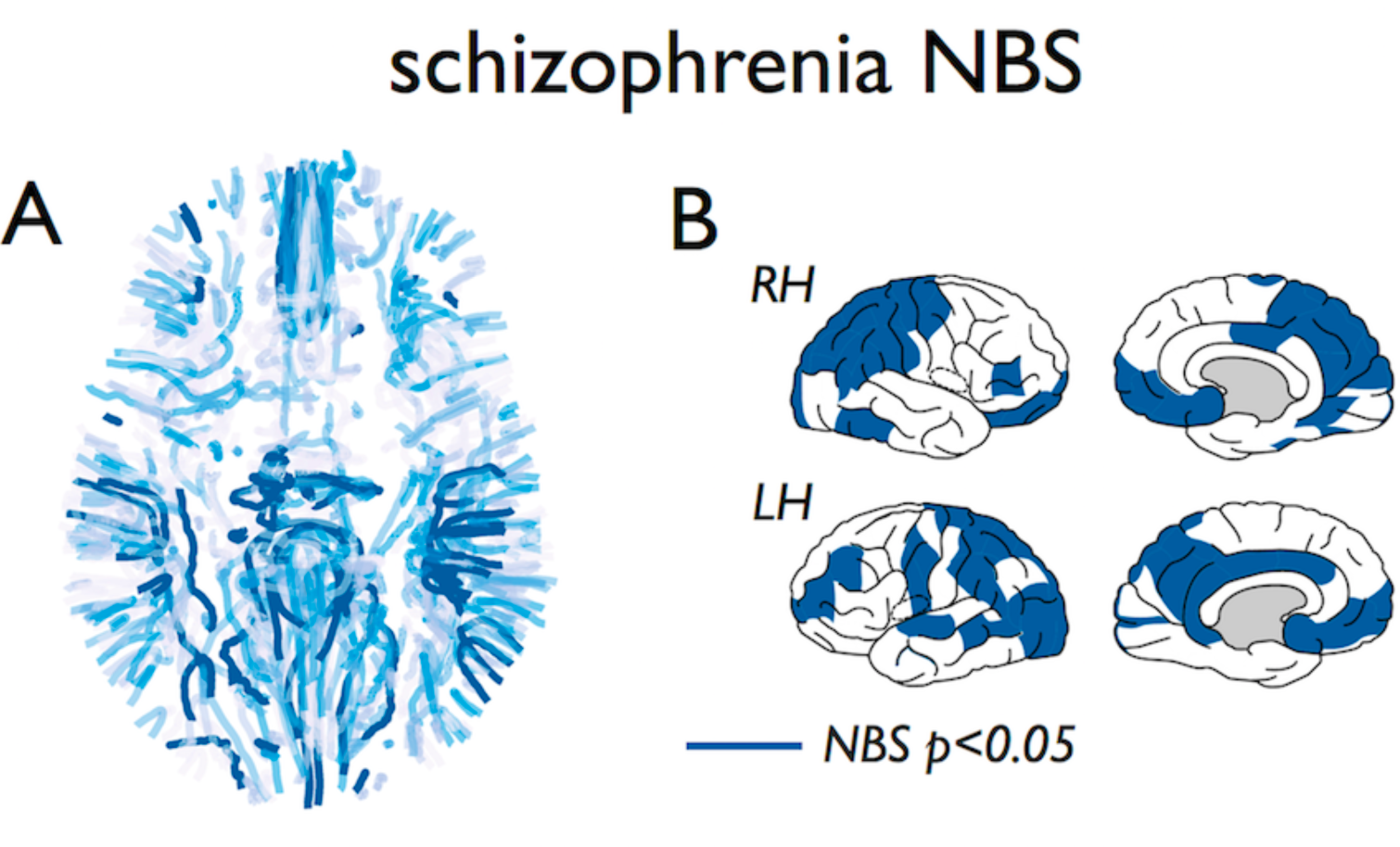
Schizophrenia disconnectivity. **Panel A** shows the cortico-cortical fiber connections of the group-average human connectome (group-threshold 60%) with level of disconnectivity as measured between n=48 schizophrenia patients and n=43 matched healthy controls, with level of disconnectivity (FA-weighted, t-score) indicated from light blue (no patient-control difference) to dark blue (patients showing significant lower FA than controls, Network Based Statistic (NBS), p=0.024). **Figure B** shows the cortical regions involved in the NBS subnetwork showing significantly reduced white matter connectivity in patients.

### Human connectome evolution and disease

We then examined the observed disease effects in schizophrenia in context of the set of human-specific connections and the set of shared connections between chimpanzees and humans. Human-specific connections (shown in Figure 3A) showed a significantly higher level of involvement in schizophrenia as compared to the set of shared connections (permutation testing with 10,000 permutations, p<0.001), in support of the hypothesis that evolutionarily novel connectivity pathways show higher involvement in disease pathology. *Robustness analysis.* Effects were replicated across two validation sets of schizophrenia patients collected from two separate patient cohort studies on schizophrenia, including n=40 patients and n=37 matched healthy controls, and n=30 and n=33 healthy controls respectively (SI Table 2). Similar to the principal dataset, both validation datasets showed uniquely human connections to display higher levels of disease involvement as compared to the set of connections shared between humans and chimpanzees (p=0.0116, p=0.0048).

**Figure 3.**
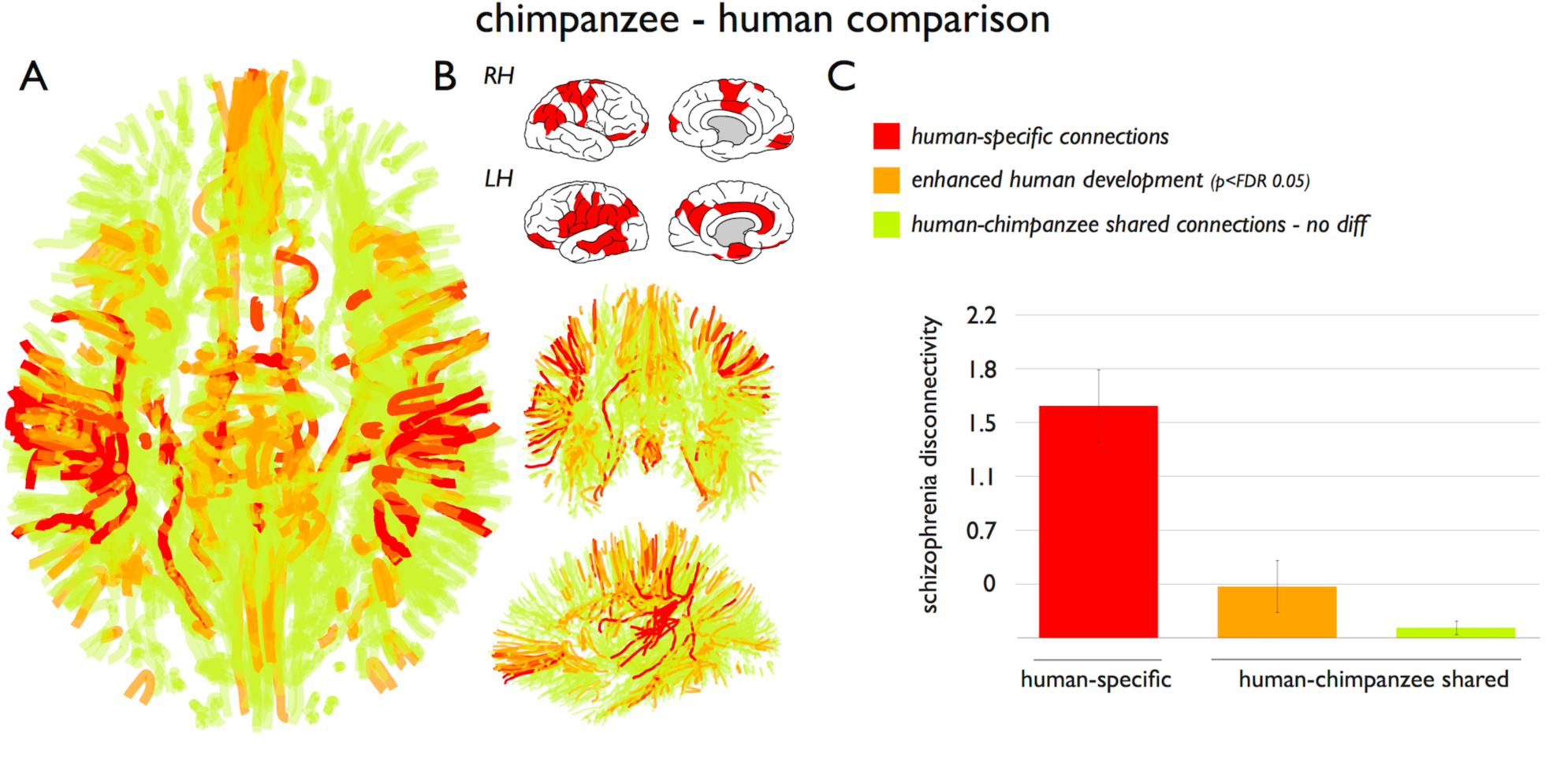
Human-specific connections and association to disease. **Panel A** identified cortico-cortical connections of the human cerebral connectome. Red connections identify human-specific connections, orange connections show human-chimpanzee shared connections showing significant positive human enhancement (normalized FA, p<0.05 FDR), green connections indicate remaining human-chimpanzee connections showing no species difference and connections of the group-average connectome. **Panel B** shows the subset of cortical regions that displayed one or more human-unique cortico-cortical connections. **Panel C** shows mean schizophrenia disease effects (y-axis: t-statistic of FA values controls versus patients) across human-specific (red) and connections shared between humans and chimpanzees. Effects in shared connections were further decomposed into connections showing enhanced human development as compared to chimpanzees (chimpanzee – human difference FDR p<0.05) and connections that showed no positive difference in normalized FA between chimpanzees and humans (p>0.05). Human-specific connections showed the largest disease effect, with human enhanced connections in second place (Jonckheere-Terpstra p<0.001).

FA was taken as the main metric of connectivity strength, a metric related to tract integrity and myelination levels (15). Studies have examined both changes in FA (e.g. (13, 14, 16)) and changes in fiber density (NOS) connectivity (e.g (17, 18)) as a metric for connectome disconnectivity in patients. We primarily focused on FA here to reflect evolutionary and disease differences in connection organization above for example number of connected streamlines or tract volume. The main reason for this is that NOS is more directly related to voxel size and brain volume, making them in our opinion less suitable for a cross-species comparison in context of disease effects (see also Supporting Results for discussion). Indeed, when we examined disease connectivity effects in terms of NOS weighted connectivity matrices no cross-species relationship between schizophrenia disease effects was observed (all p>0.05). We thus further focused on FA-based evolutionary and disease disconnectivity, arguably mostly reflecting changes in fiber structure and integrity and tract myelination.

### Human and chimpanzee shared connections

We further examined the set of shared connections between chimpanzees and humans in more detail, to examine the level of disease association across shared connections, and to rule out the possibility that differences in scan quality between species prevented us from detecting pathways in chimpanzees that were found in humans. Across the total set of connections shared between humans and chimpanzees (i.e. connections found in at least 60% of both populations) the level of evolutionary difference in connectivity strength was assessed as the t-score between the edge-wise FA values of the group of chimpanzees and the group of humans. The level of connectivity difference between chimpanzees and humans was positively correlated with the level of schizophrenia disconnectivity (r=0.17, p=0.0023). Similar effects were found in the two disease validation datasets: r=0.22, p=0.001 and r=0.17 p=0.0342), suggesting that connections showing largest differentiation between chimpanzees and humans show the largest involvement in the disorder.

We divided shared connections into two categories based on their level of evolutionary difference between chimpanzee and humans: *I* no significant difference in connection strength between chimpanzee and human and *II* enhanced human development (FDR p<0.05). Across the resulting three categories (i.e. including human-specific connections) the human-specific connections displayed the highest level of disease disconnectivity with the human-enhanced, shared connections between chimpanzees and humans in second place (p=0.0020, Jonckheere-Terpstra test, Figure 3C). This effect was replicated in the validation datasets (p=0.0040, p=0.010).

### EBB-38 atlas

We mainly used the Desikan-Killiany atlas to be consistent with previous reported brain network findings in schizophrenia (e.g. (13, 14, 16)). The DK atlas is however designed to map the gyri and sulci patterning of the human cortical mantle and a disadvantage of the use of this atlas in a human-primate neuroimaging study is thus that it does not necessarily describe homologous cytoarchitectonic regions across the two species. This may limit the comparative connectome analysis and for this reason we further validated our findings by means of a cortical atlas designed to describe homologous regions across humans and chimpanzees. We implemented the Von Economo and Bonin&Bailey cortical atlas (EBB-38), a cytoarchitectonic atlas that describes 38 homologous cortical areas (per hemisphere) across humans and chimpanzees (see Supporting Figure 3 and Supporting Materials for a description and implementation of this atlas). Analyzing the connectome data using the EBB-38 atlas revealed similar results (Figure 4): First, comparing the EBB-38 human and EBB-38 chimpanzee group connectome showed 35 human-specific tracts (tracts observed in >60% of the humans, not in the chimpanzees). These tracts were found between frontal, temporal and parietal associative cortex, including FB, FBA, FDP, TE1, TE2, TB, PH, TG, LC1, LC2 (Supporting Table 5). Second, human-specific tracts showed significantly higher levels of schizophrenia disconnectivity as compared to human-chimpanzee shared connections (p=0.0018). Similar effects were found for the two validation datasets analyzed with the EBB-38 atlas (p=0.0296, p=0.0212). Third, across the set of EBB-38 connections shared between humans and chimpanzees (i.e. connections found in at least 60% of both populations, 255 connections) the level of evolutionary difference in connectivity strength was again found to be positively associated with the level of schizophrenia disconnectivity (r=0.24; p=0.0023; replication datasets: r=0.18; p=0.0116 and r=0.17; p=0.0197).

**Figure 4.**
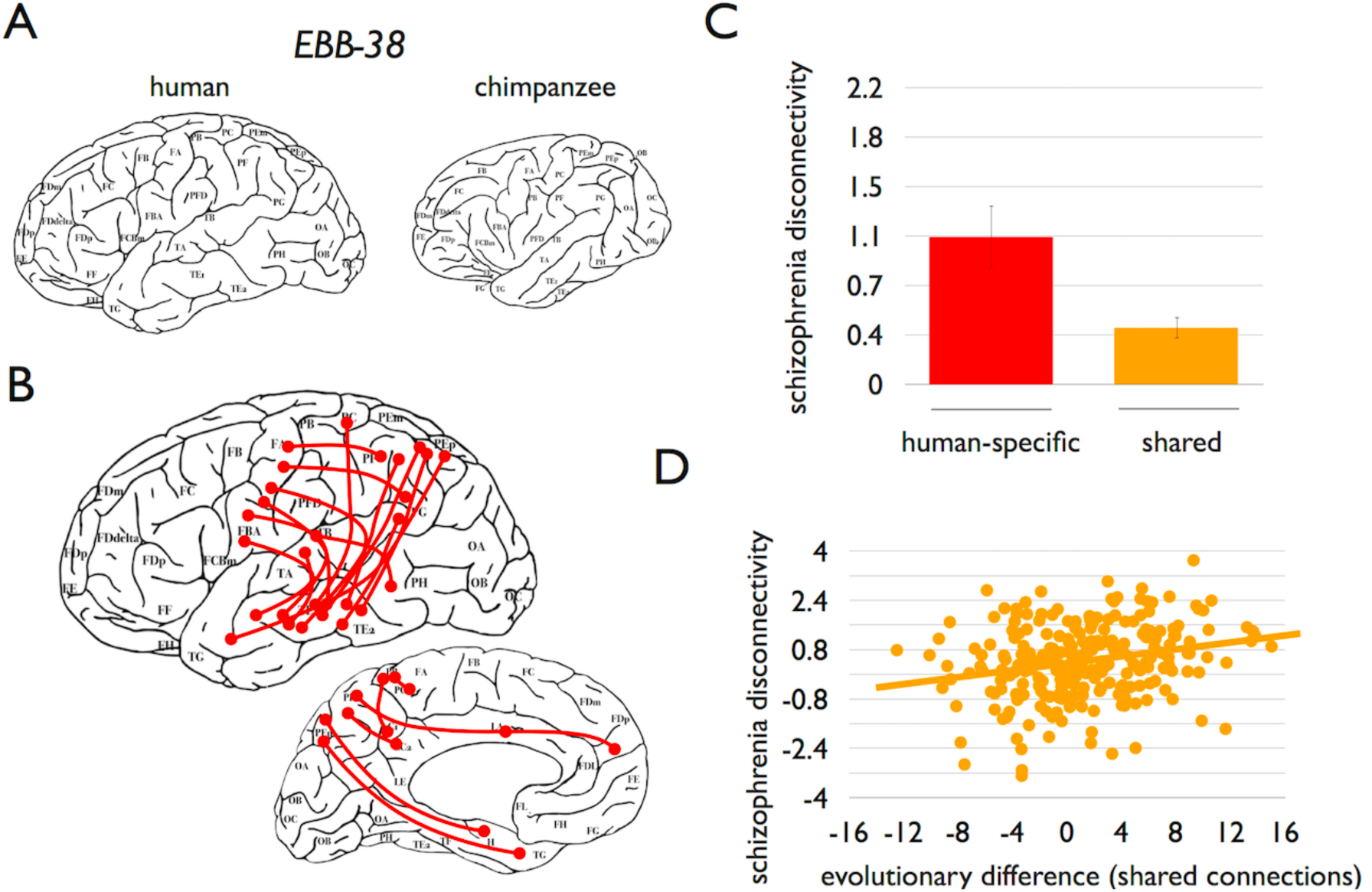
EBB-38 analysis. Figure shows results of reconstructing the human and chimpanzee connectome using the Von Economo – Bonin&Bailey human-chimpanzee homologous cortical brain atlas. **Panel A** shows the EBB-38 atlas for the human (left) and chimpanzee brain (right, see also Supplemental Materials and Supplemental Figure 3 for a detailed description of the EBB-38 atlas). **Panel B** shows a schematic representation of the EBB-38 human-specific connections drawn on the EBB-38 human atlas. Confirming findings of the main analysis, **panel C** shows larger disease involvement for human-specific tracts, as compared to human-chimpanzee shared connections. Within the set of shared connections, **panel D** shows larger disease involvement (y-axis, between-group difference t-score of tract FA) of cortico-cortical connections showing a larger evolutionary difference between human and chimpanzees (x-axis, t-score of normalized FA).

### Human-Chimpanzee-Macaque comparison

To investigate whether the human-specific connections would represent human-unique connections and with that reflect human evolutionary specializations, we further compared human and chimpanzee connections with connections in rhesus macaques, a more distantly related primate species. Macaque brains are approximately four times smaller than chimpanzee brains, and chimpanzee brains are about four times smaller than human brains (8). Looking across the three species, *human-specific connections* were again selected as the set of connections observed in humans and not in the chimpanzees and macaques, including 17, potentially human-specific, cortico-cortical connections. Similarly, human-chimpanzee-macaque shared connections were taken as the set of network connections that were reported across all three species (303 pathways, DK-114). Again, the set of connections uniquely found in humans showed significantly higher disease involvement as compared to the set of shared species connections (permutation testing, p<0.001). Similar effects were observed in the schizophrenia replication datasets (p=0.031, p=0.0020).

To contrast, we also examined the class of chimpanzee-specific connections as compared to the group of macaques, with these connections selected as the class of cortical connections observed in the population of chimpanzees (and humans) but not in the group of rhesus macaques, reflecting putative evolutionary changes in brain wiring from macaques to chimpanzees. This examination further supported specificity of the observed disease effects to human evolutionary specializations: In contrast to the class of human-specific connections, chimpanzee-specific connections (n=66) did not show higher schizophrenia-related disease effects as compared to chimpanzee-macaque shared (n=574) connections (p=0.74). In addition, again in support of a link between human evolutionary specializations and schizophrenia disease effects, no relationship was found between evolutionary changes in normalized FA in chimpanzee-macaque shared connections (i.e. between-group difference in FA in connections shared between chimpanzee and macaques) and the pattern of schizophrenia disconnectivity (r= 0.06, p=0.29).

## Discussion

Our findings suggest that evolutionary changes to cortico-cortical connections of the human cerebrum are involved in the pattern of cortical disease disconnectivity in schizophrenia. Compared to our closest living relative the chimpanzee, connections present in only humans showed on average a higher involvement in disease pathology than the majority class of connections that are shared between the two species. Furthermore, of the majority of white matter pathways shared by chimpanzees and humans, cortico-cortical connections that most differed between humans and chimpanzees were over-represented in the disease pattern of schizophrenia. Post-hoc analyses by comparing cerebral connectome organization across macaque, chimpanzee and humans further suggest that the observed disease effects are likely related to features of connectivity that evolved in the human lineage after it separated from the lineage leading to chimpanzees, that is, to human evolutionary specializations. In other words, our findings suggest that uniquely or distinctively human features of cerebral connectivity are preferentially affected in schizophrenic connectomes.

Cross-species comparisons of connectome layout over a wide range of taxa ranging from nematode to avian to mammalian species have revealed strong common principles of connectome organization (10). These putative ‘architectural principles of wiring’ tend to involve a general trade-off between a drive to minimize the expenditure of neural wiring favoring the formation of short-range local circuitry and on the other hand forces that favor the development of long-range connections and topological network attributes that are beneficial for global communication and integration (19), a network property arguably important for integrated brain function. Such a trade-off may involve increased selection pressure for long-range associative connections to keep the overall network connected in expanding brains (9) and could potentially have resulted in an increased vulnerability of relatively sparse long-range connections that manifests as disease pathology and associated disrupted cognitive functioning (2, 20). Neuroimaging studies have indeed suggested that topologically important association pathways may be particularly involved in schizophrenia (21). Taken together, our findings now suggest that the evolutionary development of such white matter pathways in the human brain to satisfy the pressure for cortico-cortical integration and with that advanced brain functionality may have contributed to pathological vulnerability for schizophrenia.

Comparative genomic examinations support an evolutionary background of schizophrenia. Evolutionary genetic markers tend to be enriched in schizophrenia (22), with schizophrenia risk having mainly developed in recent human evolution (23). A recent direct comparison of the human genome to that of chimpanzees showed schizophrenia associated loci to be enriched in genes close to human accelerated regions of the genome (24), suggesting that schizophrenia may be related to evolved specializations of the human brain. Notably, the cortical pattern of expression of schizophrenia risk genes has been suggested to resemble the pattern of disconnectivity across cortical regions in schizophrenia (25), making future study of the relationship between expression patterns of risk genes, human genomic changes and patterns of cortical connectivity an important avenue to further elucidate the underpinnings of the disorder.

In the present study, human-specific connections particularly involved cortico-cortical connections of known areas of the language network, including the posterior part of the left superior temporal sulcus, left middle temporal gyrus, left pars opercularis, left inferior parietal cortex and posterior parts of the left middle temporal gyrus. These regions have an integral role in complex language processing (e.g. (26)), a trait often argued to play a role in the disease pathology of schizophrenia. We note that these evolutionarily novel cortico-cortical connections do not necessary imply the evolution of completely new white matter tracts in humans. Instead, they could involve enhancement of evolutionarily conserved tracts between cortical areas, but with more elaborate and widespread projections resulting in new region-to-region connections in the service of enhanced brain functionality. Indeed, homologues of frontal-temporal language areas (i.e., Wernicke’s and Broca’s areas) have been reported across several primate species, but with humans exhibiting modified microstructural organization and more widespread connectivity, presumably related to development of human-unique language processing skills (27). The left middle temporal and inferior parietal cortex are also involved in advanced tool use, so the novel human connections in these areas could be involved in this function (a feature often argued in the context of human intelligence) (28) and interestingly shown to exhibit abnormal asymmetry and function in schizophrenia (e.g. (29)). Furthermore, human-specific connections also include connections of the orbitofrontal cortex, parietal and temporal associative cortex, cortical areas and connections known to play an important role in social cognition (30), emotional processing (31) and affiliative behavior (32). Deficits in social cognition are often regarded as one of the core components of schizophrenia (33), which supports the notion that the evolutionary development of social cognition may play an important role in the origin of the disorder (5).

Comparative analysis across macaque, chimpanzee and human suggests that disease disconnectivity was most severe for tracts observed uniquely in the human brain, and therewith supports thoughts on the disorder to potentially relate to human evolutionary specializations (5, 34). This may implicate that the ‘origin’ of dysconnectivity effects as observed in schizophrenia may relate to the time period after the human-chimp common ancestor. The lack of observed disease effects in macaque-chimpanzee shared connections may thus potentially suggest that previous primate species did not yet develop a connectome layout vulnerable to brain disconnectivity related to schizophrenia-like behavior. These observations are, however, somewhat contradictory with arguments of psychotic-like behavior to be evident and hypothetically possible in extant chimpanzees (5). With schizophrenia characterized as a complex psychiatric brain disorder with multiple symptom dimensions, it remains an open question to what extent the observed dysconnectivity patterns in human-unique tracts contribute to the core of the disorder and/or relate to specific symptom dimensions.

We believe that it is important to further note that from our findings it remains unknown whether the vulnerability reported here is specific to schizophrenia, or whether it describes a more general vulnerability to mental disorders. Certain psychiatric traits for example, such as low mood, depression and anxiety are found in a range of primate species (35, 36), but several other psychiatric and neurological disorders such as autism, bipolar disorder, and Alzheimer’s disease are, similar to schizophrenia, thought to be unique to humans (37, 38). Like schizophrenia, these disorders involve widespread changes in brain connectivity and evolutionary pressure on network organization may similarly have played a role in the neurobiological architecture of these disorders. Indeed, the notion that potential non-specificity of evolutionary changes in cortical connectivity may play a role in a wide range of mental health disorders is supported by GWAS studies showing significant overlap in genetic risk between schizophrenia, bipolar disorder and major depression (39).

The field of comparative connectomics is currently limited by the relatively sparse availability of connectome data from different primate species and by the variety of employed methodologies (e.g. tract-tracing versus MRI) and experimental conditions (e.g. anatomical versus functional measurements, *ex-vivo* versus *in-vivo* experiments) used to derive connectomes. To overcome these issues as much as possible, we included DWI data from a rather uniquely large sample of chimpanzees (n=22), data that was carefully acquired under as much the same conditions as possible (e.g. all *in-vivo* acquired MRI data avoiding a less-optimal comparison between post-mortem acquired animal and *in-vivo* acquired human data, matching levels of SNR across human and animal datasets, preserved cross-species voxel resolution). However, it of course remains difficult to rule out any effects of brain volume or global data differences. We therefore performed post-hoc analyses to make sure that effects could not simply be explained by global differences in FA (see SI) and/or long-range connections being more difficult to trace in chimpanzees as compared to humans due to differences in e.g. brain volume, scan quality, etcetera (see SI).

A second limitation is that our findings are inherently limited by the nature of the methodology employed. Diffusion MRI may perhaps be the only methodology currently available to measure chimpanzee and human anatomical brain connectivity *in-vivo* in relatively large numbers, but it has well-documented limitations, including difficulty reconstructing complex fiber orientations, an inability to resolve fiber direction and a recognized underestimation of both long-range and short-range connectivity. In a post-hoc analysis we verified that there was no general (i.e. non-disease related) tendency to find random between-group effects in human-specific connections (see SI). We further note that given the mentioned limitations of diffusion MRI, despite our rather strict selection of human-specific connections, we cannot fully rule out that some of the connections classified as human-specific could exist in the chimpanzee population. These tracts could have a much lower microstructural organization in chimpanzees (e.g. lower axonal count, less myelination together resulting in low DWI signal), making them not detectable in this group. Nevertheless, if that were the case, we reason that the observed effects reflect large enhancement of these cortico-cortical connections in humans, with our data suggesting that this evolutionary development covaries with high involvement in psychosis.

Taken together, our cross-species chimpanzee – human connectome comparison suggests that novel and enhanced human connections are compromised in schizophrenia, and therefore that schizophrenia may have resulted from the evolution of novel circuitry in the human brain. Evolution of the human connectome in service of developing more complex brain functionality may have come at a price of a higher risk for brain dysfunction.

## Materials and Methods

### Subjects and data acquisition

#### Primates and human dataset

MRI data were collected from 22 adult female chimpanzees (Pan trolodytes, 29.4 ± 12.8 years), 58 adult female human subjects (Homo sapiens, 42.5 ± 9.8 years) and 8 female rhesus macaques (Macaca mulatta, 14 ± 6.7 yrs). For all three species, MRI data was acquired on Siemens 3T Trio Tim Scanners and included the acquisition of a structural T1 scan and a diffusion MRI scan. We refer to (40) and Supporting Materials for a detailed description of the scanning parameters.

#### Schizophrenia dataset

MRI data of the schizophrenia dataset included the examination of 3T DWI data of 48 patients with schizophrenia and 43 age and gender matched healthy controls (demographics listed in Supporting Table 2). All participants provided informed consent as approved by the local medical ethics committee for research in humans (METC), with procedures carried out according to the directives of the “declaration of Helsinki” (amendment of Edinburgh, 2000). MRI data was acquired on a clinical Philips MRI scanner and included the acquisition of a T1 scan and a DWI scan (see Supporting Materials for scanning parameters). Effects were validated in two additional schizophrenia validation datasets, including an independent replication dataset of 40 patients and 37 healthy controls and a second independent dataset of 30 patients and 33 healthy controls (see Supporting Table 3 for demographics and details).

### Macroscale DWI connectome construction

For each individual dataset (i.e. for both primates and humans, and across all datasets), an individual connectome map was constructed by combining the cortical segmentation map with the DWI data. Each subject’s T1 image was processed using FreeSurfer, including automatic grey and white matter segmentation, followed by parcellation of the cortex using a 114 area subdivision of the Desikan-Killiany atlas of cortical structures [see Supporting Materials for other the use of other atlases]. Using the DWI images, the diffusion signal within each voxel was reconstructed and deterministic fiber tracking was used to reconstruct cortico-cortical connections (see Supporting Materials for details). A connectivity matrix of size NxN was then reconstructed by determining for each pair of cortical regions whether these two regions were interconnected by streamlines (see Supporting Materials for details). A chimpanzee and human example is shown in Supporting Figure 2. Connectivity strength of region to region pathways was taken as the average FA of the streamlines between *i* and *j,* weighted by the physical length of a streamline traveling through each voxel. FA was used as our primary metric of connectivity strength, a metric related to tract integrity and myelination levels (15) and often used to examine case-control differences in connectivity strength (14, 16).

### Schizophrenia disease disconnectivity maps

Schizophrenia disease disconnectivity was assessed by performing edge-wise comparison of the FA-weights of each reconstructed connection in the patients and controls. Between-group difference was taken as the t-statistic of a two sample t-test comparison between edge-wise FA values of the patients and controls, with higher t-statistic indicating greater disconnectivity of this particular edge in the group of patients (see SI and Results for details).

### Comparative connectomics

#### Human - chimpanzee comparison

Human-unique connections were selected as the set of connections that were observed in ≥60% of the human subjects and in none of the chimpanzee connectome maps (i.e. 0% of the chimpanzee subjects). Species shared connections were selected as the subset of cortico-cortical connections that were consistently observed in both species (≥60% of humans and ≥60% of chimpanzees, see SI for other thresholds, see also SI for HCP validation). Per shared connection, the level of species difference was computed as the between-group t-statistic between the FA values of the constructed connections of the group of chimpanzees and the group of humans (see SI for further details).

## Acknowledgements

**General:** We thank Alessandra Griffa for interesting discussions

**Funding:** MPvdHeuvel was funded by a VIDI grant from the Netherlands Organization for Scientific Research (NWO) and a Fellowship of MQ. MRI data was partly provided by the Human Connectome Project, WU-Minn Consortium (Principal Investigators: David Van Essen and Kamil Ugurbil; 1U54MH091657) funded by the 16 NIH Institutes and Centers that support the NIH Blueprint for Neuroscience Research; and by the McDonnell Center for Systems Neuroscience at Washington University.

## Author Contributions

MPvdH and JKR designed the study, analyzed the data and wrote the manuscript. LS, SL, RP analyzed the data and wrote the manuscript. MPvdH, NEMH, IS, WC, LI, TP and JR collected the data and wrote the manuscript.

## Extended and Supplemental Materials, Methods and Results

### Subjects and data acquisition

#### Primates and human dataset

MRI data were collected from 22 adult female chimpanzees (Pan trolodytes, 29.4 ± 12.8 years), 58 adult female human subjects (Homo sapiens, 42.5 ± 9.8 years) and 8 female rhesus macaques (Macaca mulatta, 14 ± 6.7 yrs). Chimpanzees and macaques were housed at Yerkes National Primate Research Center (YNPRC) in Atlanta, Georgia. Procedures were carried out in accordance with protocols approved by the YNPRC and the Emory University Institutional Animal Care and Use Committee (IACUC, approval #:YER-2001206). All humans were recruited as healthy volunteers with no known neurological conditions (IRB approval #:IRB00000028).

Primate and human data were taken from the previous study of (18) that compared hub regions across old world monkeys, apes and humans. In brief, the following procedures were followed: Macaques and chimpanzees were immobilized with ketamine (2-6 mg/kg) prior to being anaesthetized with isoflurane (1%). For all three species, MRI data was acquired on two Siemens 3T Trio Tim Scanners and included the acquisition of a structural T1 scan and a diffusion MRI scan. We used a phase-array human knee coil for macaques. For chimpanzees, we used a standard circularly polarized (CP) birdcage coil because their protruding jaw was too large to fit into the standard phase-array coil designed for humans. For humans, we used a 12-channel phase-array coil. T1 parameters included, *macaque*: MPRAGE, voxel-size: 0.5 mm isotropic, TR/TE: 2600/3.37, scan-time 25 minutes; *chimpanzee*: MPRAGE, voxel-size: 0.8 mm isotropic, TR/TE: 2600/3.06 ms, scan-time 16 minutes; *human*: MPRAGE, voxel-size: 1 mm isotropic, TR/TE: 2600/3.02 ms, scan-time 4 minutes. Diffusion MRI included the acquisition of two equal sets of diffusion weighted images with the second set acquired with a reversed phase-encoding (left-right) for removing susceptibility distortions in the post-processing (41). DWI acquisition parameters included: *macaque*: spin echo EPI, voxel-size: 1.1 mm isotropic, TR/TE: 7000/108 ms, diffusion directions: 60, b-weighting: 1000 s/mm^2^, b0-scans: 50, scan-time 86 minutes; *chimpanzee*: spin echo EPI, voxel-size: 1.8 mm isotropic, TR/TE: 5900/86 ms, diffusion directions: 60, b-weighting: 1000 s/mm^2^, b0-scans: 40, scan-time 60 minutes; *human*: spin echo EPI, voxel-size: 2 mm isotropic, TR/TE: 8500/95 ms, diffusion directions: 60, b-weighting: 1000 s/mm^2^, b0-scans: 8, scan-time 20 minutes. Diffusion MRI datasets were corrected for eddy-current, motion and susceptibility distortions (41, 42). Head motion parameters were extracted from the realignment of the diffusion images with the b0. With the chimpanzees and macaques sedated during scanning and their head and body firmly padded, they had less head movement than the (non-sedated) human subjects (p<0.001, p=0.27). We verified that this did not have an effect on our analysis, with a post-hoc analysis of matching humans and chimpanzees on head motion showing highly similar results (data shown in Supplemental Results). Acquisition protocols were designed to yield similar SNR levels of the diffusion weighted images across species (see: (18)).

#### Schizophrenia dataset

MRI data of the schizophrenia dataset included the examination of DWI data of 48 patients with schizophrenia and 43 age and gender matched healthy controls. This included an MR dataset described and examined previously in context of affected macroscale connectivity in schizophrenia (18) (demographics listed in Supplemental Table 2). Patients and controls were matched for age and gender and were part of a large ongoing cohort study. All participants provided informed consent as approved by the medical ethics committee for research in humans (METC) of the University Medical Center Utrecht, The Netherlands, with procedures carried out according to the directives of the “declaration of Helsinki” (amendment of Edinburgh, 2000). Diagnostic consensus of patients was achieved in the presence of a psychiatrist according to DSM-IV criteria for schizophrenia. All patients were receiving typical or atypical antipsychotic medication at time of scanning (see Supplemental Table 2 for details).

MRI data was acquired on a clinical Philips MRI scanner and included the acquisition of a T1 scan and a DWI scan. T1 acquisition parameters included: 3D FFE using parallel imaging, 0.75 mm isotropic, TR/TE: 10/4.6ms, 200 slices covering whole head. DWI acquisition parameters included: spin echo EPI, two equal sets with the second set acquired with a reversed phase-encoding direction, voxel size: 2.0 mm isotropic, TR/TE: 7035/68 ms, diffusion directions: 30, b-weighting: 1000 s/mm^2^, b0-scans: 5, scan-time 12 minutes. T1 scans were processed with FreeSurfer, including segmentation of grey and white matter and parcellation of the cortex into 114 cortical areas using a subparcellation of the Desikan-Killiany atlas (43, 44). Pre-processing of the DWI data included correction for eddy-current, motion and susceptibility distortions (41, 42) and realignment of the cortical segmentation map with the b0. There was no difference in motion between the patient and control datasets (p=0.29).

### Schizophrenia replication datasets

Effects were validated in two additional schizophrenia datasets, including an independent replication dataset of 40 patients and 37 healthy controls and a second independent dataset of 30 patients and 33 healthy controls. Demographics of these additional datasets are listed in Supplemental Table 3. Both replication datasets included the same DWI and T1 data acquisition as the principal schizophrenia dataset acquired on a similar type of 3T Philips clinical scanner as the principal dataset. There was again no difference in motion between the patient and control datasets (p=0.16, p=0.81). The principal and two replication datasets were included as separate datasets in this study as the three cohorts differed in the time period of data acquisition (principal dataset: year 2011-2013; replication dataset 1: 2014-2016; replication dataset: 2015-2017), the topic of investigation (respectively: brain morphological changes, neural substrate of hallucinations, neural substrate of self-agency) and distribution of demographics (see Supplemental Table 2 and 3).

### Macroscale DWI connectome construction

For each individual dataset (i.e. for both primates and humans, and across all datasets), an individual connectome map was constructed by combining the cortical segmentation map with the DWI data. First, a subject’s T1 image was processed using FreeSurfer (45), including automatic grey and white matter segmentation, followed by parcellation of the cortex using a 114 area subdivision of the Desikan-Killiany atlas of cortical structures (43, 44) [Other atlas including respectively a coarser parcellation with 68 cortical regions (DK-68), a more fine-grained parcellation with 219 cortical regions (DK-219) revealed similar findings, see results]. Segmentations were manually checked and corrected where needed. Supplemental Figure 1 shows segmentations of exemplary human, chimpanzee and macaque subjects.

Second, using the DWI images, the diffusion signal within each voxel was reconstructed using combined tensor reconstruction and Compressed Sensing for DWI (schizophrenia dataset)(46) or Q-sampling imaging (human - primate dataset) (47), allowing for the reconstruction of complex diffusion fiber configurations (i.e. crossing/kissing fibers), with in case of no indication of a complex fiber model the main diffusion direction represented by a single tensor. Within each white matter voxel, the level of *fractional anisotropy* (FA) was computed as a metric of preferred diffusion direction using the fitted tensor model, and interpreted as a metric of pathway microstructural organization (14, 15, 48). Third, deterministic fiber tracking was used to reconstruct cortico-cortical connections. Within each white matter voxel, eight evenly distributed seeds were placed and streamlines were started from each of the eight seeds following the best matching diffusion direction from voxel to voxel until a streamline reached one of the stopping criteria (i.e. reached the gray matter, exited the brain mask, made a turn of >60 degrees and/or reached a voxel with low fractional anisotropy <0.05; other stop settings (i.e. max degree > 45, FA<0.1) resulted in similar findings). Fourth, combining the collection of constructed streamlines and the cortical segmentation map, a connectivity matrix of size NxN was reconstructed by determining for each pair of N cortical regions whether these two regions were interconnected by one or more streamlines. Setting a fiber threshold (i.e. taking only connections that included over *t* fibers) (12, 49) to reduce the inclusion of potential false positive streamline reconstructions in the connectivity matrix did not change the nature of our findings (data shown in Supplemental Results). Supplemental Figure 2 shows the reconstructed fiber bundles of an exemplary chimpanzee and human dataset.

Connectivity strength of region to region pathways was taken as the average FA of the streamlines between *i* and *j.* For each streamline in the selected pathway, the FA values of the set of voxels that were crossed by the streamline were selected and the average FA of a streamline was computed by taking the mean over this set of voxels. With a streamline describing a 3D pathway traveling through the MRI volume, crossing each voxel differently, the mean was taken across the set of voxels taking into account the proportion that each streamline spend in each of the crossed voxels (50) (51). Next, obtained streamline FA values were averaged over the included streamlines to get the average FA value of the connection between region *i* and region *j*, which was then stored in the FA weighted connectivity matrix. In this combined disease - comparative examination we chose FA as a metric of connectivity strength, a metric related to tract integrity and myelination levels (15, 52) and often used to examine case-control differences in connectivity strength (4, 13, 14, 16).

### FA vs NOS as a metric of connectivity strength

In this combined disease - comparative examination we chose FA as a metric of connectivity strength, a metric related to tract integrity and myelination levels (15) and often used to examine case-control differences in connectivity strength (4, 16, 53). We note in this context that we preferred FA to reflect evolutionary differences in connection organization above for example number of connected streamlines (NOS) or tract volume (e.g. (18, 50)), as NOS is directly related to voxel size and brain volume. There was no relationship between global normalized FA and number of streamlines included in the connectome reconstruction, in neither the group of chimpanzees (p=0.802) nor human samples (p=0.28), suggesting no general difference in global normalized FA that could be attributed to differences in brain size. As expected, there was also no significant difference in global normalized FA between the group of chimpanzees and humans (p=0.4099). Studies (including of our own group) have taken *both* changes in FA (e.g. (14) (13) (16)) *and* changes in NOS connectivity (e.g (17, 18)) as a metric for connectome disconnectivity in patients. Taken together, for our cross-species disease study, we believe that using NOS as a metric of evolutionary and disease differences would thus be a less preferred metric for comparison to changes related to schizophrenia. In our opinion, FA is a more sensible metric in our combined cross-species - disease study. As mentioned in the main text, we explicitly note that we indeed did not find any cross-species relationship in NOS and schizophrenia disease effects (all p>0.05). It was for this reason that we focused on FA as a metric of evolutionary differences and disease disconnectivity, taken as a metric related to tract integrity and myelination levels (15).

### Schizophrenia disease disconnectivity map

Across all FA connectivity matrices, a schizophrenia disease disconnectivity map was formed by performing edge-wise comparison of the FA-weights of each reconstructed connection in the patients and controls. For each connection observed in at least 50% of the patient and 50% of the control population, between-group difference was taken as the t-statistic of a two sample t-test comparison between edge-wise FA values of the patients and controls, with higher t-statistic indicating greater disconnectivity of this particular edge in the group of patients. Within this edge-wise evaluation, per edge, missing values (i.e. of a subject in which no information on FA of that connection was present due to a non-existing connection) were excluded from the edge-wise t-test. As a side effect of this statistical approach the degrees-of-freedom (*df*) can vary across edges, due to a varying number of data points per evaluated network edge. We verified that there was no significant association between edge prevalence (i.e. the number of times an edge was found to be present in a group of subjects) and random between-group t-statistic (p=0.4469). Finally, using other group prevalence thresholds (e.g. 40%, 60% or 70%) revealed similar results (data shown in Supplemental Results). The same statistical procedures were performed for the two schizophrenia validation datasets.

### Comparative connectomics

#### Human - chimpanzee comparison

Human-unique connections were selected as the set of connections that were consistently observed in the human connectome maps (≥60% of the human subjects) and in none of the chimpanzee connectome maps (i.e. 0% of the chimpanzee subjects). Species shared connections were selected as the subset of cortico-cortical connections that were consistently observed in both species (≥60% of humans and ≥60% of chimpanzees).

A prevalence group threshold of 60% (human) and an exclusion threshold of 0% (chimpanzee) were used to reduce the inclusion of false-positive reconstructions at the group level (11) and thus to include a rather strict definition of human-specific tracts. To reduce the potential inclusion of false positive tracts (11) and to use a strict definition of human-specific and human-chimpanzee shared connections (12) other connections (i.e. tracts observed in less than 60% of the groups and of which one can not be sure whether they represent true positives) were not considered for further analysis. Other prevalence thresholds revealed similar findings (see Supplemental Results). Using Human Connectome Project data for human connectome reconstruction also revealed similar findings (Supplemental Results).

For each of the cortico-cortical pathways in the subset of shared connections, the level of species difference was computed as the between-group t-statistic between the FA values of the constructed connections of the group of chimpanzees and the group of humans. FA-weighted connectivity matrices were redistributed to a normalized distribution with equal mean and standard deviation (0.5, 0.15 respectively) before comparison to allow for human-chimpanzee cross-species comparison of connection weights irrespective of potential global differences in FA. Per connection shared across the two species, a t-test across the values of the chimpanzee and human specimens was performed, with the level of difference in connectivity strength between the chimpanzees and humans then taken as the resulting t-statistic.

#### Human-Chimpanzee-Macaque comparison

To investigate whether the human-specific connections would represent human-unique connections and therewith human evolutionary specializations, we further compared human and chimpanzee connections with connections in rhesus macaques, a more distantly related primate species.

Macaque brains are approximately four times smaller than chimpanzee brains, and chimpanzee brains are about four times smaller than human brains (8, 54–57). Looking across the three species, *human-specific connections* were again selected as the set of connections observed in humans (more than 60% of the group) and not in the chimpanzees (i.e. 0%) and macaques (i.e. 0%), including 17, potentially human-specific, cortico-cortical connections. Similarly, human-chimpanzee-macaque shared connections were taken as the set of network connections that were reported across all three species (i.e. in over 60% of all 4 populations, 303 pathways, DK-114 atlas).

### EBB-38 human chimpanzee homologues atlas analysis

#### Human-chimpanzee homologous atlas

In the main analysis, the Desikan-Killiany atlas was used to parcellate the cortex into 114 distinct cortical regions. This atlas was used for the patient-control analysis as well as the comparative analysis. We choose to use the DK atlas to provide consistency with previous reports on schizophrenia disconnectivity. However, with the Desikan-Killiany atlas built for a gyri-based parcellation of the human cortex, a disadvantage of this approach is that this parcellation does not describe homologous regions across the chimpanzee and human brain. With atlas selection known to have an effect on network reconstruction (e.g. (44)) this could potentially influence our presented comparative connectome results. To rule this out and to overcome this methodological limitation we also analyzed the comparative data (and the patient and control data) with the Von Economo – Bonin&Bailey (EBB-38) human and chimpanzee brain atlas, a cortical parcellation atlas that describes homologous cortical areas across the two species. To this end, we created a species specific FreeSurfer version of the EBB-38 atlas for both human and chimpanzee, describing 38 homologous cytoarchitecturally defined regions per hemisphere in the human and chimpanzee brain. This atlas was based on a combination of the 1950 chimpanzee cortical atlas of Von Bonin and Bailey (58) and the 1925 human cortical atlas of Von Economo and Koskinas (59). Below we describe the FreeSurfer implementation of the human and chimpanzee version of this atlas.

#### Human EBB-38

We earlier built a FreeSurfer version of the Von Economo - Koskinas atlas for the human brain (60). This was performed by manually segmenting the cytoarchitectonically defined cortical areas as defined by Von Economo and Koskinas onto a template human brain, from which a FreeSurfer atlas file was built using the FreeSurfer atlas building tools. A detailed description of the segmentation and atlas build procedures are given in (60), which also includes a public version of this atlas. Here, we re-used this 3D Von Economo-Koskinas FreeSurfer atlas as the basis to construct the human cortical version of the human-chimpanzee homologous EBB-38 atlas. Out of the 43 human cortical parcels in our 2016 FreeSurfer version of the Von Economo-Koskinas 1925 atlas, the cytoarchitectonic cortical areas as defined by Von Bonin and Bailey for the chimpanzee brain were constructed (see below for the description of these regions). This was performed on the basis of the writings of Von Bonin and Bailey in their 1950 report (58) in which they in high detail describe 44 cortical areas for the chimpanzee brain, and how these areas relate to the human brain. In particular, they provide a detailed description of how the Von Economo human cortical areas relate to the coarser mapping observed in the chimpanzee cortex. In their parcellation of the chimpanzee cortex (see description of how we made the chimpanzee atlas in the next paragraph), Bonin and Bailey specifically described that they observed less cytoarchitectural variation in most areas of the cortex, meaning fewer further subdivisions of cortical areas in the chimpanzee brain as compared to the reports of Von Economo for the human brain. For example, in the cingulate cortex of the chimpanzee brain Von Bonin and Bailey described one anterior cingulate region LA and described that they could not find a further subdivision of this region in the chimpanzee cortex, while Von Economo and Koskinas could further split this larger area LA into smaller subregions LA1 and LA2 based on cytoarchitectonic differences in the human brain. The smallest subdivisions as described in *both* the chimpanzee by Von Bonin and Bailey and the human by Von Economo and Koskinas were included in the EBB-38 human-chimpanzee homologous atlas. Supplemental Table 4 summarizes the mapping of these 38 regions to Von Economo – Koskinas areas of the human cortex (as described in (60, 61)) and their relationship to the 38 Von Economo – Bonin&Bailey (EBB-38) human-chimpanzee homologous areas.

Next, the resulting human version of the EBB-38 atlas was run for each of the human datasets included in this study, including the human subjects in the comparative analysis and the patients and controls of the three included schizophrenia datasets. The human EBB-38 atlas is shown in Supplemental Figure 3A; Supplemental Figure 3C shows the application of this atlas to an exemplary human subject of the dataset.

#### Chimpanzee EBB-38

A similar procedure was used to build the by Von Bonin and Bailey described chimpanzee version of the EBB-38 atlas. As mentioned, Von Bonin and Bailey examined and described the cytoarchitectonic division of the chimpanzee cortex using the same nonmenclature of cortical areas as used by Von Economo and Koskinas in their 1925 writing. As such, Von Bonin and Bailey defined the cortical areas of the chimpanzee cortex in close comparison with the cytoarchitectural regions described by Von Economo for the human brain. In every lobe of the chimpanzee cortex they compared and described in detail whether and how the same divisions could be made based on the same cytoarchitectural characteristics (such as cortical layer thickness, neuronal cell type, size and density) similar to those in the human brain.

We note that, in total, Von Bonin and Bailey originally described 44 cortical regions for the chimpanzee brain, 38 of which we found possible to include in the FreeSurfer parcellation. Out of the 44 we noted that 6 areas were too narrow or located in a sulcus (being regions: PCop, FCop, FDT, PA, PD) or embedded within another cortical region (TB), and we could therefore not properly segment them and include them in the FreeSurfer atlas segmentation. This resulted in the described total of 38 cortical regions per hemisphere.

We refer to this atlas as the EBB-38 (*Economo-Bonin&Bailey 38 region)* atlas. Following the FreeSurfer guidelines, for all defined regions in the EBB-38 atlas, region borders were plotted onto a FreeSurfer reconstructed chimpanzee cortical surface to create individual region label files, now specific to the chimpanzee brain. Next, as performed for the reconstruction of the human version (see above), a colortable describing a unique RGB color scheme for all labels was combined with the label files to form a chimpanzee EBB-38 FreeSurfer *annotation* file, one per hemisphere. Left and right hemisphere annotation files were then used for FreeSurfer atlas training, forming one chimpanzee specific EBB-38 atlas file per hemisphere. The resulting chimpanzee EBB-38 FreeSurfer atlas was then applied to each of the 22 chimpanzee subjects, providing individual cortical EBB-38 segmentations. The chimpanzee EBB-38 atlas is shown in Supplemental Figure 3B; Supplemental Figure 3D shows the application of this atlas to an exemplary chimpanzee subject of the dataset.

#### EBB-38 connectome reconstruction and group analysis

Next, the derived human and chimpanzee EBB-38 cortical atlas was used to reconstruct EBB-38 connectome maps for each of the datasets used in this study. For this, the same procedures were used as applied for the reconstruction of the connectome maps based on the DK-114 atlas. Per subject, the set of reconstructed streamlines was combined with the EBB-38 cortical segmentation map and for each subject a connectivity matrix of size 38×38 was reconstructed by determining for each pair of cortical regions whether regions were interconnected by one or more streamlines. Connectivity strength of each reconstructed cortical pathway was taken as the average FA of the streamlines between cortical regions. This resulted in new EBB-38 connectome maps for both the comparative analysis (i.e. human and chimpanzee connectome maps) and for the three schizophrenia datasets.

Next, based on the new EBB-38 connectome maps, human-specific connections were selected as those cortico-cortical connections that could be observed in more than 60% of the humans and in none of the chimpanzees. Human-chimpanzee shared connections were similarly selected as those connections observed in at least 60% of both populations.

Next, schizophrenia effects were computed for the EBB-38 connectomes. Similar as to the analysis of the DK-114 analysis, edge-wise FA values were compared by means of t-testing between patients and controls, resulting in an edge-wise disease disconnectivity map. Next, the comparative analyses were compared to the schizophrenia results, with disease effects compared between human-specific connections and human-chimpanzee shared connections. Finally, similar to the DK-114 analysis, disease effects were correlated to evolutionary differences in normalized FA within the class of human-chimpanzee shared connections. Results of these EBB-38 based analyses are represented in the Results section of the main text, and confirmed our main findings of human-specific connections to be more involved in schizophrenia disconnectivity compared to human-chimpanzee shared connections.

### Supplemental Results

#### Human-Chimpanzee-Macaque comparison

The set of connections uniquely found in humans showed again significantly higher disease involvement as compared to the set of shared species connections (permutation testing, p<0.001). Similar effects were observed in the schizophrenia replication datasets (p=0.031, p=0.0020). To contrast, we also examined the class of chimpanzee-specific connections as compared to the group of macaques, with these connections selected as the class of cortical connections observed in the population of chimpanzees (and humans) but not in the group of rhesus macaques, reflecting putative evolutionary changes in brain wiring from macaques to chimpanzees. This examination further supported specificity of the observed disease effects to human evolutionary specializations: In contrast to the class of human-specific connections, chimpanzee-specific connections (n=66) did not show higher schizophrenia-related disease effects as compared to chimpanzee-macaque shared (n=574) connections (p=0.74). In addition, again in support of a link between human evolutionary specializations and schizophrenia disease effects, no relationship was found between evolutionary changes in normalized FA in chimpanzee-macaque shared connections (i.e. between-group difference in FA in connections shared between chimpanzee and macaques) and the pattern of schizophrenia disconnectivity (r= 0.06, p=0.29).

#### Group threshold validation

In the comparative analysis in the main text human-specific tracts were selected as connections present in 60% of the human population and in 0% of the chimpanzee population (indicated by T=60/0%) and human-chimpanzee shared connections were selected as connections present in at least 60% of both populations. These thresholds were used to reduce the influence of potential false positive connection reconstructions in the analysis (11, 12, 60). In a post-hoc analysis we verified that other group selection thresholds revealed similar findings. Using a T=50/0% selection threshold revealed similar findings: human-specific tracts (n=38) showed higher schizophrenia disconnectivity effects as compared to the set of shared connections (n=579)(p<0.001) and within the set of shared human-chimpanzee connections evolutionary difference was positively correlated to disease disconnectivity (r=0.14, p=0.0035). Using a T=70/0% selection threshold revealed similar findings: human-specific tracts (n=19) showed higher schizophrenia disconnectivity effects as compared to the set of shared connections (n=322)(p<0.001) and within the set of shared human-chimpanzee connections evolutionary difference was positively correlated to disease disconnectivity (r=0.17, p=0.0025).

Also loosening the strict threshold of 0% revealed similar findings: Using a T=60/20% selection threshold showed human-specific tracts (n=91) to display higher schizophrenia disconnectivity effects as compared to the set of shared connections (n=428)(p<0.001) and within the set of shared human-chimpanzee connections evolutionary difference was positively correlated to disease disconnectivity (r=0.16, p=0.0028).

We also verified the settings of the group thresholding in the EBB-38 atlas analysis. A group threshold of 50% or 70% revealed similar findings as the main EBB-38 analysis. EBB-38 human-specific tracts (50%: n=46; 70%: n=24) showed higher schizophrenia disconnectivity as compared to the set of EBB-38 shared connections (n=331; n=202)(p=0.0177; p=0.0337) and within the set of EBB-38 shared human-chimpanzee connections, evolutionary difference was positively correlated to disease disconnectivity (r=0.24, p=0.0092; r=0.23, p=0.0011).

#### Fiber threshold validation

In the main comparative analysis cortico-cortical connections were selected when there was at least one streamline present between cortical areas. This selection criterion was used to result in a strict criterion for the selection of human-specific tracts: this selection criterion allowed for a liberal tract selection in the chimpanzees, i.e. one streamline would already result in a potential chimpanzee connection and therewith a strict human-specific selection criterion in combination with the 60/0% group selection criterion (see above). We note in this context that our main value of examination was the average FA over these tracts, not the number of reconstructed streamlines, which has been argued to not be very sensitive to the underlying number of streamlines (62). Indeed, also in this examination no clear relationship was found between global tract FA and number of reconstructed streamlines per subject (reported in main text).

However, as an alternative selection criterion, studies have advocated the use of a fiber selection threshold *fT* to reduce potential false-positive reconstructions in the individual connectome maps (12, 62). We verified that our findings were consistent when using other connection selection criteria. Using a fiber selection criterion of *fT* >2 (i.e. including connections that consisted of 3 or more streamlines) revealed similar findings: human-specific tracts (n=26) showed higher schizophrenia disconnectivity effects as compared to the set of shared connections (n=395)(p<0.001) and within the set of shared human-chimpanzee connections evolutionary difference was positively correlated to disease disconnectivity (r=0.16, p=0.0032). Also the use of a rather strict fiber selection criterion of *fT* >5 revealed similar findings: human-specific tracts (n=25) showed higher schizophrenia disconnectivity effects as compared to the set of shared connections (n=357)(p<0.001) and within the set of shared human-chimpanzee connections evolutionary difference was positively correlated to disease disconnectivity (r=0.17, p=0.0024).

We also verified the use of *fT* thresholding in the EBB-38 atlas analysis. The use of a fiber selection criterion of *fT* >5 revealed similar findings as the main EBB-38 analysis. EBB-38 human-specific tracts (n=25) showed higher schizophrenia disconnectivity as compared to the set of EBB-38 shared connections (n=95)(p=0.0260). In addition, within the set of EBB-38 shared human-chimpanzee connections, evolutionary difference again showed a positive correlation to disease disconnectivity (r=23, p=0.0357).

#### DK-68 and DK-219 validation

In the main analysis of our study we used a 114 cortical area subparcellation (44) of the Desikan-Killiany (DK) cortical atlas (43). The DK atlas is designed to identify the sulci/gyri patterning of the human brain (43) (see for a discussion on this in the context of comparative human-chimpanzee analysis the main text). With atlas resolution shown to have an effect on connectome reconstruction and case-control graph analysis (e.g. (44, 63, 64)), we verified that our findings were consistent for other DK resolutions. In a post-hoc analysis, findings were validated using the DK-68 (dividing each hemisphere in 34 cortical areas) and the DK-219 atlas (defining 111 left hemisphere and 108 right hemisphere regions). Both alternative atlases revealed findings similar to the DK-114 atlas.

Human-specific tracts (DK-68 n=10; DK-219 n=178) were found to display larger disease effects as compared to human-chimpanzee shared connections (DK-68 n=136; DK-219 n=1024) in both the DK-68 (p=0.0240) and DK-219 condition (p<0.001). Overlapping the results shown for the DK-114 atlas, examining disease effects within the class of shared connections showed a positive association between evolutionary human-chimpanzee difference in tract FA and schizophrenia disconnectivity (DK-68: r=0.19, p=0.0246; DK-219: r=0.17, p=0.0025).

#### EBB-38 atlas

Analyzing the connectome data using the EBB-38 atlas revealed similar results (Figure 4): First, comparing the EBB-38 human and EBB-38 chimpanzee group connectome showed 25 human-specific tracts (tracts observed in >60% of the humans, not in the chimpanzees). These tracts were found between frontal, temporal and parietal associative cortex, including FB, FBA, FDP, FDL, TE1, TE2, TB, PH, TG, LC1, LC2 (Supplemental Table 5). Second, human-specific tracts showed significantly higher levels of schizophrenia disconnectivity as compared to human-chimpanzee shared connections (p=0.0059). Similar effects were found for the two validation datasets analyzed with the EBB-38 atlas (p=0.0290, p=0.0036). Third, across the set of EBB-38 connections shared between humans and chimpanzees (i.e. connections found in at least 60% of both populations, 280 connections) the level of evolutionary difference in connectivity strength was again found to be positively associated with the level of schizophrenia disconnectivity (r=0.21; p=0.0023; replication datasets: r=0.27; p=0.0048 and r=0.17; p=0.0368).

#### HCP validation

In the main comparative analysis we used chimpanzee and human data acquired on the same clinical scanner for comparison. To match SNR across the two datasets, voxel-resolution was different between the chimpanzee and human datasets (chimpanzee: 1.8 mm isotropic, human: 2.0 mm isotropic). However, with higher resolution (1.25 mm) data of the human brain available through means of the Human Connectome Project (65, 66) we also verified our comparative findings with this dataset.

To this end, we analyzed the high-resolution HARDI DWI data of 487 subjects of the Human Connectome Project (HCP, Q4 release, voxel-size 1.25 mm isotropic, TR/TE 5520/89.5 ms, three runs of each 90 diffusion directions with diffusion weighting 1000, 2000 and/or 3000 s/mm^2^). Data was preprocessed using FSL (https://fsl.fmrib.ox.ac.uk/fsl/) including correction for eddy current and susceptibility distortions (see (65) for details). Similar to the analysis performed for the 3T human data in out main study, for each HCP subject tissue classification and DK-114 cortical parcellation was performed on basis of a high-resolution T1 anatomical image (voxel size: 0.7 mm isotropic). Next, based on the preprocessed HCP DWI data, white matter pathways were reconstructed using combined tensor and generalized Q-sampling imaging (GQI), allowing for the reconstruction of complex diffusion fiber configurations (i.e. crossing/kissing fibers), and streamline tractography (see (47) (25)). Eight streamlines were started in each white matter voxel, following the most matching diffusion direction from voxel to voxel until a streamline reached the gray matter, exited the brain tissue, made a turn of >45 degrees or reached a voxel with a low fractional anisotropy (<0.1) (see for details on connectome reconstruction also (67)). Next, combining the DK-114 cortical parcellation and DWI data, a weighted structural connectivity matrix of size NxN was formed, selecting from the total collection of reconstructed tractography streamlines those that touched both regions *i* and *j,* for all pairs (*i*, *j*). Tract averaged FA was used as a weight of the reconstructed cortico-cortical network connections. Similar to the chimpanzee data (and the human data in the main analysis), FA weights were resampled to a Gaussian distribution (mean:0.5, std:0.15) (50) to allow for cross-species comparison (see also main text). Finally, combining the reconstructed connectome matrices of the 487 HCP subjects, a group-averaged structural connectivity matrix was formed by taking the non-zero mean of the individual matrices, including a connection between brain regions when it was found in at least 60% of the total group of HCP subjects.

After HCP connectome reconstruction we re-performed our comparative analysis using the HCP connectome maps (instead of our own human data). To this end, similar as in the main analysis, human-specific connections were selected when cortico-cortical connections were found to be present in at least 60% of the human population (now 487 HCP subjects) and in 0% of the chimpanzee group; human-chimpanzee shared connections were selected as tracts observed in at least 60% of both populations. This analysis revealed similar findings as in our main analysis: human-specific tracts (n=31) showed larger disease effects as compared to human-chimpanzee shared connections (p=0.0038) and within the class of shared connections (n=321) a positive association was observed between evolutionary difference in tract FA between chimpanzee and human and schizophrenia disconnectivity (r=0.14, p=0.033).

#### Head motion analysis

The chimpanzee subjects were sedated and their head and body firmly padded during scanning, resulting in significantly less head motion as compared to the human population (p<0.001). To verify that this did not have an effect on our comparative analysis, we repeated our analysis with a subset of the humans which showed as a group no difference in motion as compared to the chimpanzee sample. From the total set of human subjects, the top 22 (the same n as the group of chimpanzees was selected) humans with the least motion were selected. This subset showed no significant difference in head motion with the group of chimpanzees (p=0.612). With this subset of human subjects the comparative analysis was repeated and data was compared to the disease effects. This analysis revealed similar findings to including the total set of human subjects. Human-specific tracts (n=25) showed higher disease effects as compared to human-chimpanzee shared connections (p=0.0044) and within the class of shared connections (n=410) a positive association was observed between evolutionary human-chimpanzee difference in tract FA and schizophrenia disconnectivity (r=0.13, p=0.013).

To further verify that head motion did not bias our methods into finding erroneous new tracts, we also contrasted set A of top 22 human subjects with the lowest movement (n=22 was selected to match the sample size of the chimpanzees) to set B of human subjects with the highest level of motion (i.e. the remaining human subjects). No difference in tracts was found between group A and B, showing that difference in head motion did not result in a bias to finding group-specific tracts.

#### Analysis of fiber length

We performed post-hoc analyses to make sure that effects could not simply be explained by long-range connections being more difficult to trace in chimpanzees as compared to humans due to differences in e.g. brain volume, scan quality, etcetera. First, human-specific connections were not the longest traced pathways (in absolute distance), with other long-range connections equally well reconstructed in both chimpanzees and humans. For each of the reconstructed uniquely human connections there were on average 24 (std:13) connections that were longer and similarly reconstructed in the group of the chimpanzees. In addition, the prevalence of the set of uniquely human connections was not significantly different from the prevalence of the set of longer connections shared across the two species (p>0.05), suggesting that the reported long-range connections could be equally well-traced across the two species.

#### Analysis on random disease effects

In a permutation analysis, patients and controls were randomly mixed and a ‘random disconnectivity map’ (1,000 randomizations examined) was formed and correlated to the described chimp-human connectivity map, which now no longer resulted in a significant difference in group effects between human-specific connections and connections shared between humans and chimpanzees (p>0.05). Furthermore, effects were also not present when an additional post-hoc analysis was performed in which FA-values were randomly mixed (1,000 times) within each individual before computation of the disease between-group map (p>0.05), suggesting that there was also no overall bias to find random disease effects in one type of connection, for example across-species different and/or long-range cortico-cortical connections.

## Supplemental Figures

**Supplementary Figure 1.**
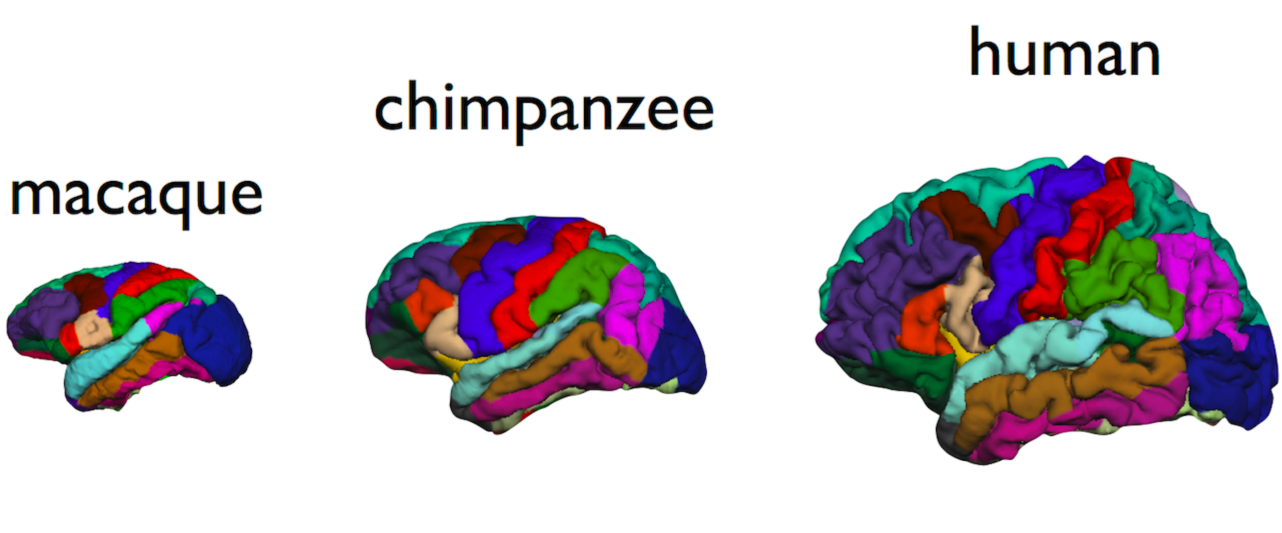
Example of cortical segmentation of an exemplary macaque (left), chimpanzee (middle) and human specimen (right).

**Supplementary Figure 2.**
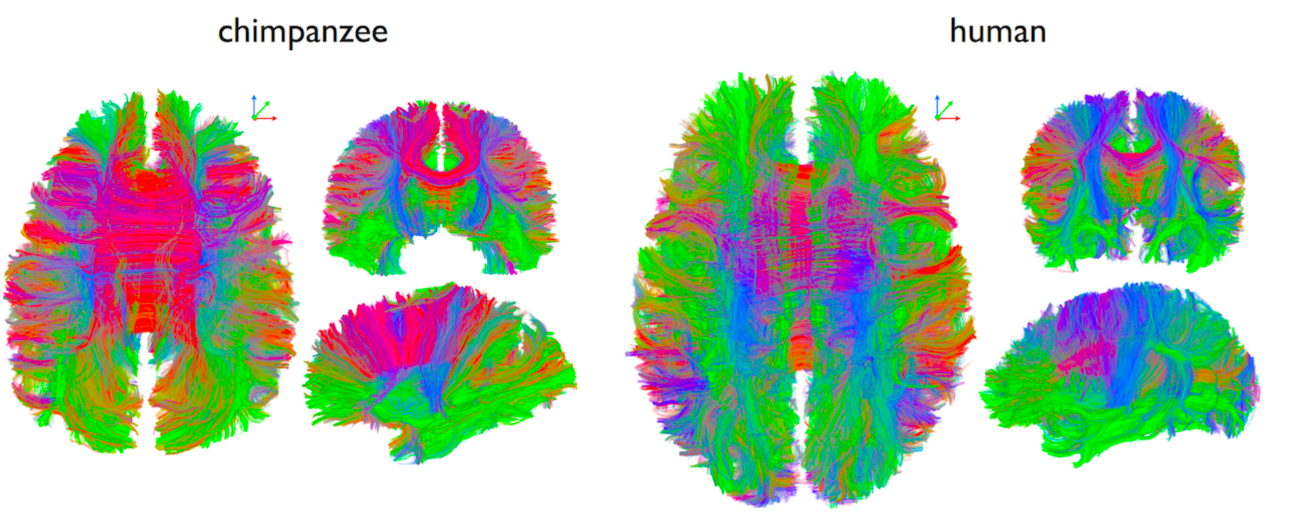
Example of connectome fibertracking of an exemplary chimpanzee (left) and human specimen (right).

**Supplementary Figure 3.**
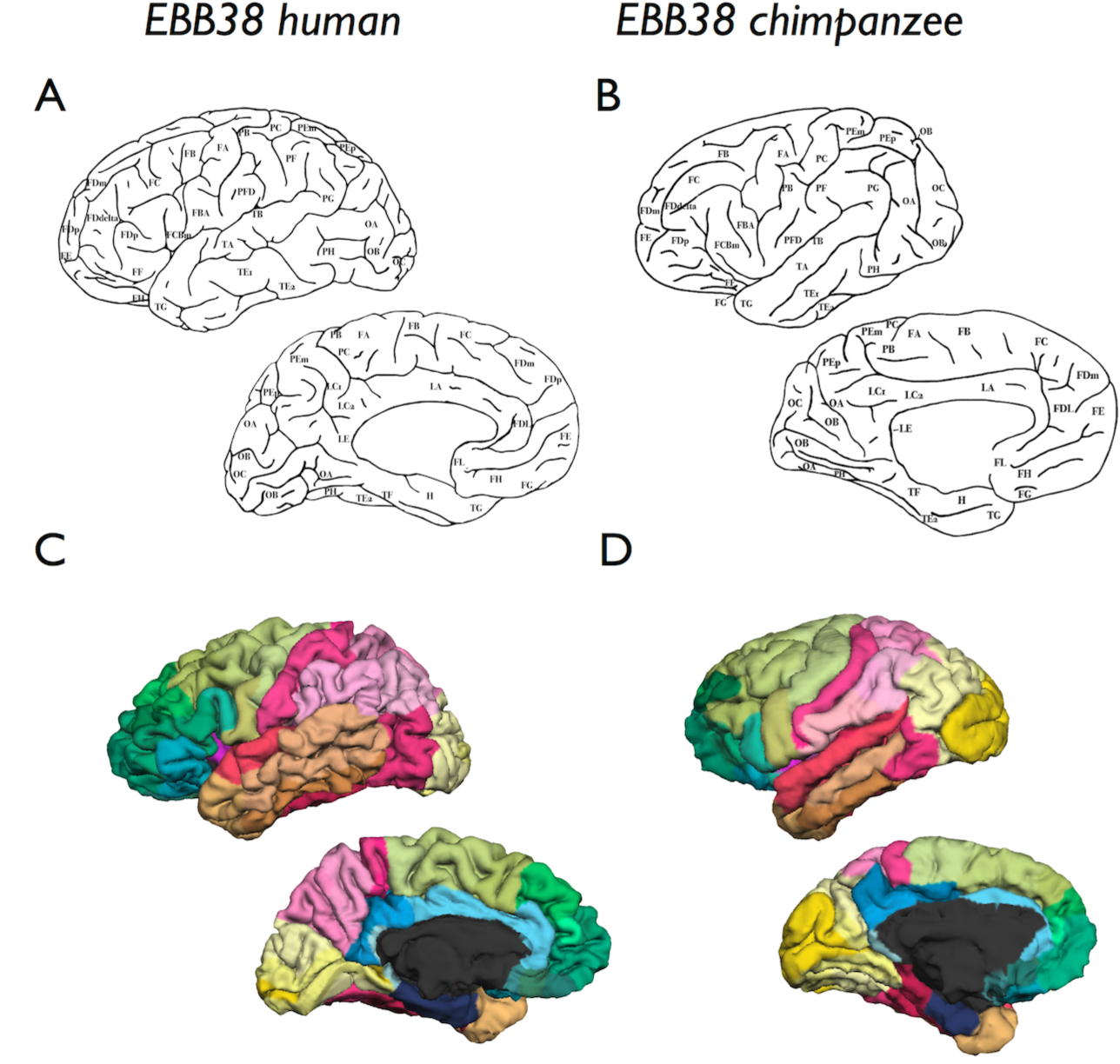
Panel **A** and **B** show the Von Economo and Bonin and Bailey 38 cytoarchitectonically defined areas of respectively the human and chimpanzee cortex. Panel **C** and **D** show the 3D application of the EBB-38 atlas to respectively an exemplary human (panel C) and chimpanzee (panel D) subject of the datasets.

**Supplementa1 Table 1.** List of observed human-specific connections.

| Cortical region of DK-114<sup>a</sup> | Cortical region of DK-114<sup>a</sup> |
| --- | --- |
| lh precentral ventral | lh bankssts |
| lh precuneus | lh caudalanteriorcingulate |
| lh posterior superiorparietal | lh entorhinal cortex |
| lh parsopercularis | lh posterior inferiorparietal |
| lh postcentral | lh posterior inferiorparietal |
| lh ventral precentral | lh posterior inferiorparietal |
| lh ventral precentral | lh posterior inferiortemporal |
| lh supramarginal gyrus | lh posterior inferiortemporal |
| rh frontalpole | lh lateralorbitofrontal |
| lh postcentral | lh anterior middletemporal |
| lh ventral precentral | lh anterior middletemporal |
| lh ventral precentral | lh anterior middletemporal |
| lh supramarginal gyrus | lh anterior middletemporal |
| lh ventral precentral | lh posterior middletemporal |
| lh supramarginal gyrus | lh posterior middletemporal |
| rh posterior superiorfrontal | lh posteriorcingulate |
| lh supramarginal gyrus | lh ventral precentral |
| lh rostralanteriorcingulate | lh precuneus |
| rh posteriorcingulate | lh precuneus |
| rh postcentral | rh anterior inferiorparietal |
| rh ventral precentral | rh anterior inferiorparietal |
| rh postcentral | rh posterior inferiorparietal |
| rh ventral precentral | rh posterior inferiorparietal |
| rh superiorparietal | rh posterior inferiorparietal |
| rh supramarginal gyrus | rh posterior inferiorparietal |
| rh parsorbitalis | rh lingual |
| rh supramarginal gyrus | rh ventral precentral |

<sup>a</sup> DK-114 regions describe the 114 cortical regions of the subparcelation of the Desikan-Kiliany atlas. lh = left hemisphere; rh=right hemisphere

**Supplementa1 Table 2.**
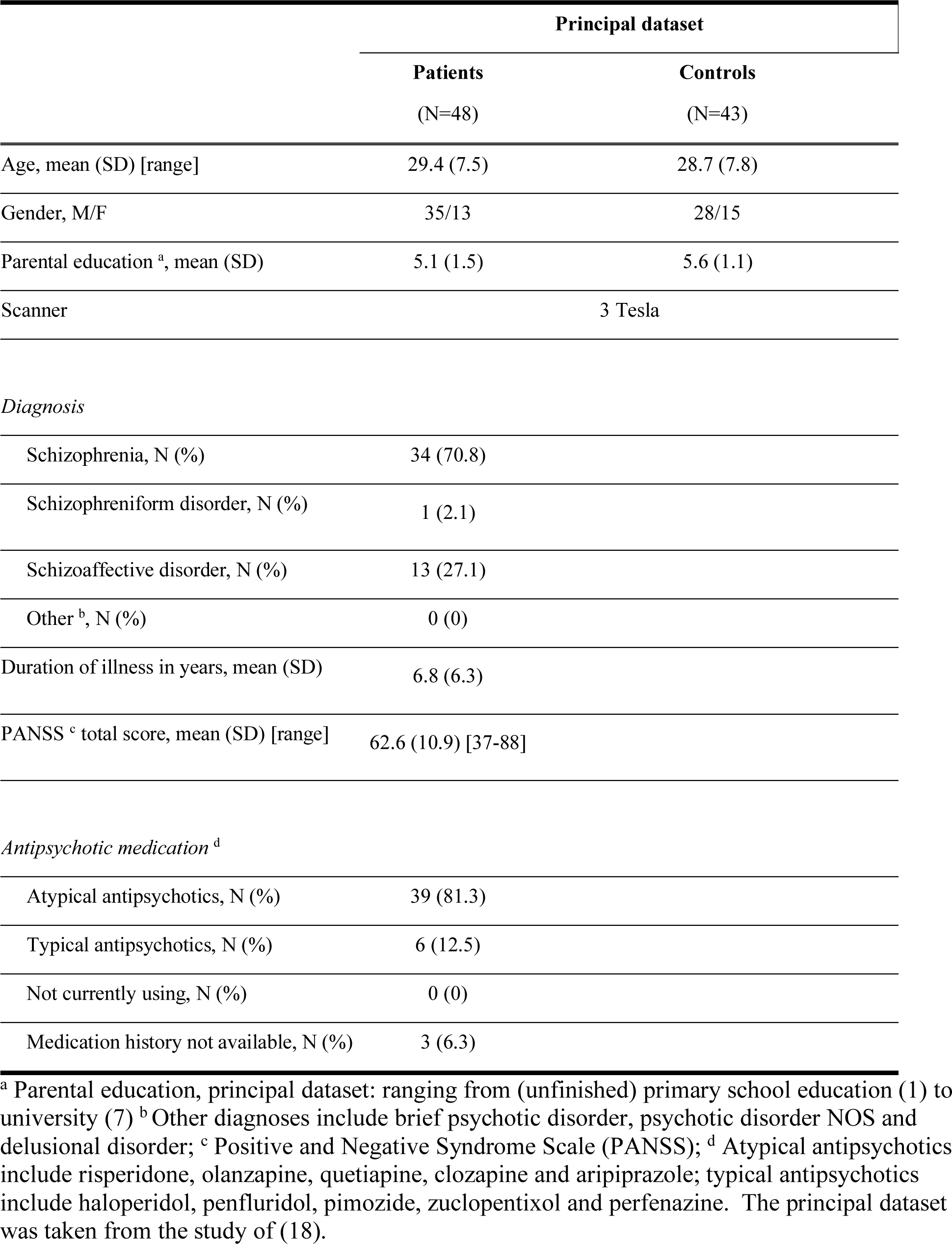
Demographic and clinical characteristics of principal dataset

|  | Principal dataset |  |
| --- | --- | --- |
|  | Patients<br>(N=48) | Controls<br>(N=43) |
| Age, mean (SD) [range] | 29.4 (7.5) | 28.7 (7.8) |
| Gender, M/F | 35/13 | 28/15 |
| Parental education <sup>a</sup>, mean (SD) | 5.1 (1.5) | 5.6 (1.1) |
| Scanner | 3 Tesla |  |

*Diagnosis*
|  |  |
| --- | --- |
| Schizophrenia, N (%) | 34 (70.8) |
| Schizophreniform disorder, N (%) | 1 (2.1) |
| Schizoaffective disorder, N (%) | 13 (27.1) |
| Other <sup>b</sup>, N (%) | 0 (0) |
| Duration of illness in years, mean (SD) | 6.8 (6.3) |
| PANSS <sup>c</sup> total score, mean (SD) [range] | 62.6 (10.9) [37-88] |

*Antipsychotic medication <sup>d</sup>*
|  |  |
| --- | --- |
| Atypical antipsychotics, N (%) | 39 (81.3) |
| Typical antipsychotics, N (%) | 6 (12.5) |
| Not currently using, N (%) | 0 (0) |
| Medication history not available, N (%) | 3 (6.3) |

<sup>a</sup> Parental education, principal dataset: ranging from (unfinished) primary school education (1) to university (7) <sup>b</sup> Other diagnoses include brief psychotic disorder, psychotic disorder NOS and delusional disorder; <sup>c</sup> Positive and Negative Syndrome Scale (PANSS); <sup>d</sup> Atypical antipsychotics include risperidone, olanzapine, quetiapine, clozapine and aripiprazole; typical antipsychotics include haloperidol, penfluridol, pimozide, zuclopentixol and perphenazine. The principal dataset was taken from the study of (18).

**Supplementa1 Table 3.** Demographic and clinical characteristics of replication datasets

|  | Replication dataset 1 |  | Replication dataset 2 |  |
| --- | --- | --- | --- | --- |
|  | Patients<br>(N=40) | Controls<br>(N=37) | Patients<br>(N=30) | Controls<br>(N=33) |
| Age, mean (SD) [range] | 37.1 (13.4) | 39.6 (14.5) | 29.5 (6.3) | 30.2 (6.8) |
| Gender, M/F | 25/15 | 10/27 | 24/6 | 26/7 |
| Parental education <sup>a</sup>, mean (SD) | 6 (2) | 4 (6) | 6 (2) | 6 (2) |

| Scanner | 3 Tesla | 3 Tesla |
| --- | --- | --- |
| *Diagnosis* |  |  |
| Schizophrenia, N (%) | 28 (70) | 29 (96.7) |
| Schizophreniform disorder, N (%) | 0 (0) | 1 (3.3) |
| Schizoaffective disorder, N (%) | 4 (10) | 0 (0) |
| Other <sup>b</sup>, N (%) | 8 (20) | 0 (0) |
| Duration of illness in years,<br>mean (SD) | 13.7 (12.4) | 9.1 (7.0) |
| PANSS <sup>c</sup> total score,<br>mean (SD) [range] <sup>#</sup> | 59.8 (14.7) [38-91] | 46.0 (8.8) [30-62] |
| *Antipsychotic medication* <sup>d</sup> |  |  |
| Atypical antipsychotics, N (%) | 22 (55) | 25 (83) |
| Typical antipsychotics, N (%) | 7 (17.5) | 3 (10) |
| Not currently using, N (%) | 7 (17.5) | 1 (3.3) |
| Medication history not available, N<br>(%) | 5 (12.5) | 1 (3.3) |

<sup>a</sup> Parental education, principal dataset: ranging from (unfinished) primary school education (1) to university (8). <sup>b</sup> Other diagnoses include brief psychotic disorder, psychotic disorder NOS and delusional disorder; <sup>c</sup> Positive and Negative Syndrome Scale (PANSS); <sup>d</sup> Atypical antipsychotics include risperidone, olanzapine, quetiapine, clozapine and aripiprazole; typical antipsychotics include haloperidol, penfluridol, pimozide, zuclopentixol and perphenazine.

**Supplementa1 Table 4.** Equivalent regions between Von Bonin&Bailey chimpanzee and Von Economo – Koskinas human atlas

| Region number | Von Bonin & Bailey – chimpanzee regions | Scholtens (2016) / Von Economo – Koskinas (1925) regions |
| --- | --- | --- |
| 1 | FA | FA |
| 2 | FB | FB |
| 3 | FBA | FBA |
| 4 | FC | FC |
| 5 | FCBm | FCBm |
| 6 | FDdelta | FDdelta |
| 7 | FDL | FDL |
| 8 | FDm | FDm |
| 9 | FDp | FDp |
| 10 | FE | FE |
| 11 | FF | FF |
| 12 | FG | FG |
| 13 | FH | FH |
| 14 | FL | FLMN |
| 15 | H | HA HB HC |
| 16 | IA | IA |
| 17 | IB | IB |
| 18 | LA | LA1 LA2 |
| 19 | LC1 | LC1 LC3 |
| 20 | LC2 | LC2 LD |
| 21 | LE | LE1 LE2 |
| 22 | OA | OA |
| 23 | OB | OB |
| 24 | OC | OC |
| 25 | PB | PB PA |
| 26 | PC | PC |
| 27 | PEm | PEm |
| 28 | PEp | PEp |
| 29 | PF | PF |
| 30 | PFD | PFD |
| 31 | PG | PG |
| 32 | PH | PH |
| 33 | TA | TA |
| 34 | TB | TB TC TD |
| 35 | TE1 | TE1 |
| 36 | TE2 | TE2 |
| 37 | TF | TF |
| 38 | TG | TG |

**Supplementa1 Table 5.**
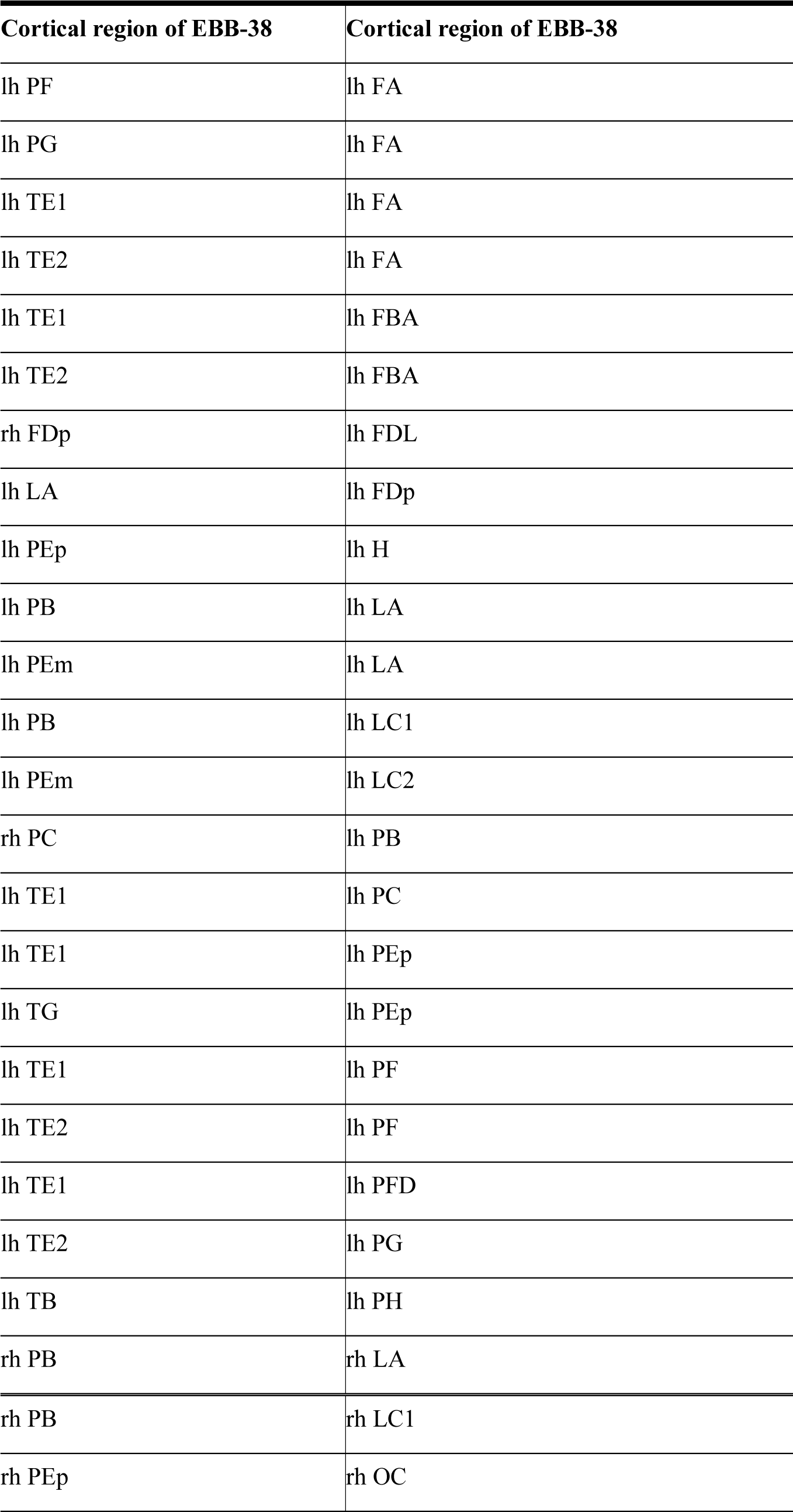
List of observed human-specific connections EBB-38 atlas.

